# MOT: a Multi-Omics Transformer for multiclass classification tumour types predictions

**DOI:** 10.1101/2022.11.14.516459

**Authors:** Mazid Abiodoun Osseni, Prudencio Tossou, Francois Laviolette, Jacques Corbeil

## Abstract

**Motivation:** Breakthroughs in high-throughput technologies and machine learning methods have enabled the shift towards multi-omics modelling as the preferred means to understand the mechanisms underlying biological processes. Machine learning enables and improves complex disease prognosis in clinical settings. However, most multi-omic studies primarily use transcriptomics and epigenomics due to their over-representation in databases and their early technical maturity compared to others omics. For complex phenotypes and mechanisms, not leveraging all the omics despite their varying degree of availability can lead to a failure to understand the underlying biological mechanisms and leads to less robust classifications and predictions.

**Results:** We proposed MOT (Multi-Omic Transformer), a deep learning based model using the transformer architecture, that discriminates complex phenotypes (herein cancer types) based on five omics data types: transcriptomics (mRNA and miRNA), epigenomics (DNA methylation), copy number variations (CNVs), and proteomics. This model achieves an F1-score of 98.37% among 33 tumour types on a test set without missing omics views and an F1-score of 96.74% on a test set with missing omics views. It also identifies the required omic type for the best prediction for each phenotype and therefore could guide clinical decisionmaking when acquiring data to confirm a diagnostic. The newly introduced model can integrate and analyze five or more omics data types even with missing omics views and can also identify the essential omics data for the tumour multiclass classification tasks. It confirms the importance of each omic view. Combined, omics views allow a better differentiation rate between most cancer diseases. Our study emphasized the importance of multi-omic data to obtain a better multiclass cancer classification.

**Availability and implementation:** MOT source code is available at https://github.com/dizam92/multiomic_predictions.

## Introduction

The development of high-throughput techniques, such as nextgeneration sequencing and mass spectometry, have generated a wide variety of omics datasets: genomics, transcriptomics, proteomics, metabolomics, lipidomics, among others. This reveals different biological facets of the clinical samples that open up new perspectives within the framework of personalized medicine. Although the majority of past studies (1–4) use a single omic data type, with a significant emphasis on genomics, transcriptomics and proteomics, there is currently a switch towards multi-omics studies. The objective is to provide a deeper and better understanding of patients’ internal states, enabling accurate clinical decision-making (5, 6). The positive impact of these multi-omics studies using machine learning techniques can already be seen in several indication areas: Central Nervous Systems (7, 8), oncology (9–12), cardiovascular diseases (13) single-cell analysis in humans (14–16). A typical multi-omics study only uses the transcriptomic data (mRNA and miRNA) and the epigenomics data (DNA methylation also known as CpG sites). However, there is a multitude of other omics data types that must be taken into consideration for a complete assessment of a patient internal state. Many reasons are often invoked for not considering other omics: heterogeneity (5), missing values, outliers and data imbalances (17). But the most important is the under-representation of certain omics types in databases due to limited effort to acquire this type of data, costs associated with their acquisition and the technical decisions made by laboratory groups. Lately, several studies (18–20) are studying cancer diseases under the prism of personalised medicine. These studies are trying to unveil the varying sources responsible for the cancer disease at a micro level i.e. for each patients. The varying sources imply that the different omics available may have various impacts on each cancer patients.

To exploit all these data, the development of computational methods has accelerated. The rapid growth and success of machine learning and deep learning models have led to an exponential increase of applications models to biological problems including the cancer diseases classification task. For instance, a traditional auto-encoder (21) was used to embed some multi-omics data (mRNA, miRNA and DNA methylation) into a 100-dimensional space to identify multi-omics features linked to the differential survival of patients with liver cancer (10). Xu et *al*. (22), introduced HI-DFN Forest, a framework built for the cancer subtype classification task. The framework includes a multi-omics data integration step based on hierarchical stacked auto-encoders (23) used to learn an embedded representation from each omics data (mRNA, miRNA and DNA methylation). The learned representations are then used to classify patients into three different cancer subtypes: invasive breast carcinoma (BRCA), glioblastoma multiform (GBM) and ovarian cancer (OV). Targeting a different perspective on the multi-omics data usage, Li et *al*. (24) addressed the task of predicting the proteome from the transcriptome. To achieve this task, Li et *al*. (24) built three models: a generic model to learn the innate correlation between mRNA and protein level, a random-forest classifier to capture how the interaction of the genes in a network control the protein level and finally a trans-tissue model, which captures the shared functional networks across BRCA and OV cancers. It should be noted that most of these studies used only one omic view to tackle the cancer identification or classification task. As for pan-cancer with multi-omics data, (25) introduced DeepProg, a semi-supervised hybrid machine-learning framework made essentially of an auto-encoder for each omics data type to create latent-space features which are then combined later to predict patient survival subtypes using a support vector machine (SVM). DeepProg is applied on two omics views (mRNA and DNA methylation) for 32 cancer types from the TCGA portal (https://www.cancer.gov/tcga). OmiVAE, (26) on the other hand, is a variational auto-encoder based model (27), used to encode different omics datasets (mRNA and DNA methylation) into a low-dimensional embedding on top of which a fully connected block is applied to the classification of the 33 tumours from UCSC Xena data portal (28). These models are limited in the number of omics and which ones, they can integrate successfully.

To respond to the lack of existing model integrating and processing many different omics views with missing views for samples, we introduce MOT, a multi-omic transformer architecture. Initially introduced to solve Sequence to Sequence (Seq2Seq) translation problems, the transformer model (29) is widely applied to various domains and is increasingly becoming one of the most frequently used deep learning models. This model includes two main parts: an encoder and a decoder composed of modules (multi-heads attention mechanisms and feed forward layers). The modules can be stacked on top of each other multiple times. The popularity of the transformer architecture lies in the attention heads mechanism that offers a level of interpretability of the model’s decision process. We perform a data augmentation step in the learning phase to obtain a robust MOT model handling missing omics data type. Data augmentation encompasses techniques used to increase the amount of data by adding altered copies of already existing data or newly created synthetic data from existing data. The impact of this method is well demonstrated in the literature (30–32). Here, new examples were created from the original samples by randomly generating alternate subsets of omics data type available for the examples. We compared the MOT performance to some baselines algorithms. To our knowledge, this is the first model that integrates and processes up to five omics data types regardless of their availability and offers a macro level of interpretability for each phenotype for the pan-cancer multiclass classification task.

## Material and Methods

### Datasets and preprocessing

#### Datasets

The TCGA pan-cancer dataset is available on the UCSC Xena data portal. There are 33 tumour types in the dataset. Five types of omics data, mRNA (RNA-Seq gene expression), miRNA, DNA methylation, copy number variation (CNVs) and protein, were used in this study. Among them, three (mRNA, DNA methylation, CNVs) are datasets of high-dimensional space. The gene expression (mRNA) profile of each sample comprises 20532 identifiers referring to corresponding genes. A log2 transformation (log2(norm_value+1)) was applied on the original count resulting in an mRNA version called the batch effects normalized mRNA. The Illumina Infinium Human Methylation BeadChip (450K) arrays provide DNA methylation profiles with 485,578 probes. The Beta value of each probe represents the methylation ratio of the corresponding CpG site. The CNVs profile of each sample comprises of 24776 identifiers which are estimated values from the ones measured experimentally. The estimated values are −2, −1, 0, 1, 2, which represent respectively homozygous deletion, single copy deletion, normal diploid copies, low-level copy number amplification, or high-level copy number amplification. As for the miRNA profile, it is comprised of 743 identifiers. The values of the miRNA dataset were also log2-transformed. Finally, the protein expression dataset is comprised of 210 identifiers. All the omics datasets were downloaded from the UCSC Xena data portal on September 1st, 2021. As most omics datasets, the dataset is imbalanced: there is a discrepancy in the availability of samples for each tumour type. It is a well-documented problem (17) specific to this kind of dataset. To illustrate this, the authors refer readers to figure 3 in supplementary data which present the number of samples available for each of the 33 tumours in the dataset. The imbalance is easily observable as as we have more than 1200 samples for breast cancer and fewer than 50 samples for cholangiocarcinoma (bile duct cancer). Table 4 in supplementary data presents all the 33 cancer types with their abbreviations.

#### Preprocessing

A feature selection step was performed on the omics datasets with a high-dimensional space to comprehensively integrate all of the omic dataset. The targeted omics datasets are the mRNA, the DNA methylation and the CNVs. The dimension reduction step, a standard step in multi-omics data processing, is well documented in many studies. For example, Wu and *al*. (33) presented many feature selections and techniques adapted to multi-omics problems. Here, we apply the median absolute deviation 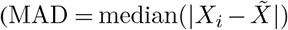 with 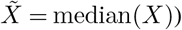 which is a robust measure of the variability of a univariate sample of quantitative data. The MAD was applied to the mRNA and the DNA methylation datasets. Regarding the CNVs dataset, it contains categorical values [(−2, −1, 0, 1, 2)]. Thus, another feature selection method was applied, the mutual_info_classif, available on sickit-learn (34), which estimates the mutual information for a discrete target variable. Mutual information (MI) between two random variables is a non-negative value, which measures the dependency between the variables. It is equal to zero if and only if two random variables are independent, and higher values mean higher dependency. Since it can be used for univariate features selection, we believed it was the most suitable for the CNVs dataset. From each applied method on the targeted omics dataset, we selected 2000 features per omics type. It should be recalled that the miRNA and proteomics dataset were used directly without a feature selection step. After the dimension reduction step, the omics dataset were integrated using the parallel integration method (33) which consists of putting together all the omics available together to obtain a matrix with *n* rows (the samples) and *m* column (the omics features). There is no consensus on the integration method in the studies but Wu and *al*. (33) presented an excellent review of all the main techniques used. As for the data augmentation step, new samples were built by randomly selecting a subset of the omics views initially available for the sample. Thereby for each patient from the original dataset built earlier, a combination between 1 and 4 views is randomly selected and replaced with 0. Amongst the five omics datasets targeted, a sample must have at least one of those omics data available to be considered in the final dataset.

### MOT: a transformer model

The transformer model is constituted of encoders and decoders and is built around the attention mechanism. Each encoder includes two principal layers: a self-attention layer and a feed-forward layer. Before feeding the input data to the encoder, the input is passed through the embedding layer which is a simple linear neuronal network. Let 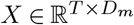 an input data consisting of *T* tokens in *D_m_* dimensions. Similar to the NLP framework where each token *t* represents a word in a sentence, the token here represents the numerical value of the multi-omic data concerned. Let’s denote 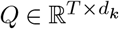, the matrix containing all query vectors of all the omic datasets, 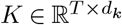, the matrix of keys and 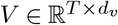, the matrix of all values. The query represents a feature vector that describes what we are looking for in the sequence. The key is also a feature vector which roughly describes what the element is “offering”, or when it might be important. The value is also a feature vector which is the one we want to average over. *T* is the length of the sequence, *d_k_* is the hidden dimension of the keys and *d_v_* the hidden dimension of the values. Thus the self attention value is obtained by:

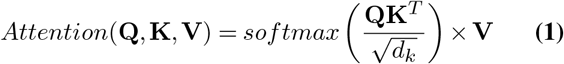

The multi-head attention is the integration of multiple single self-attention mechanism to focus simultaneously on different aspects of the inputs. Literally it represents a concatenation of single head attention mechanism. The initial inputs to the multi-head attention are split into *h* parts, each having queries, keys, and values. The multi-head attention is computed as follows:

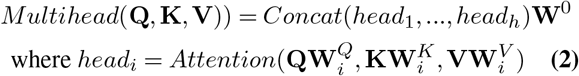

with 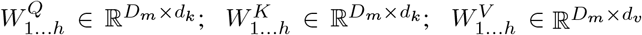 and 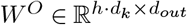. The attention weights are then sent to the decoder block which objective is to retrieve information from the encoded representation. The architecture is quite similar to the encoder, except that the decoder contains two multi-head attention submodules instead of one in each identical repeating module. In the original transformer model, due to the intrinsic nature of the self-attention operation which is permutation invariant, it was important to use proper positional encoding to provide order information to the model. Therefore, a positional encoding step 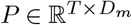 was added after the embedding step. Here, in our multi-omic task, the order of the inputs is not important since there is no relation between the features. Therefore, our multi-head attention layers do not include the positional encoding module. Figure 1 illustrates the MOT model which is the original model introduced by Vaswani et *al*. (29) without the positional encoding step.

**Fig. 1.**
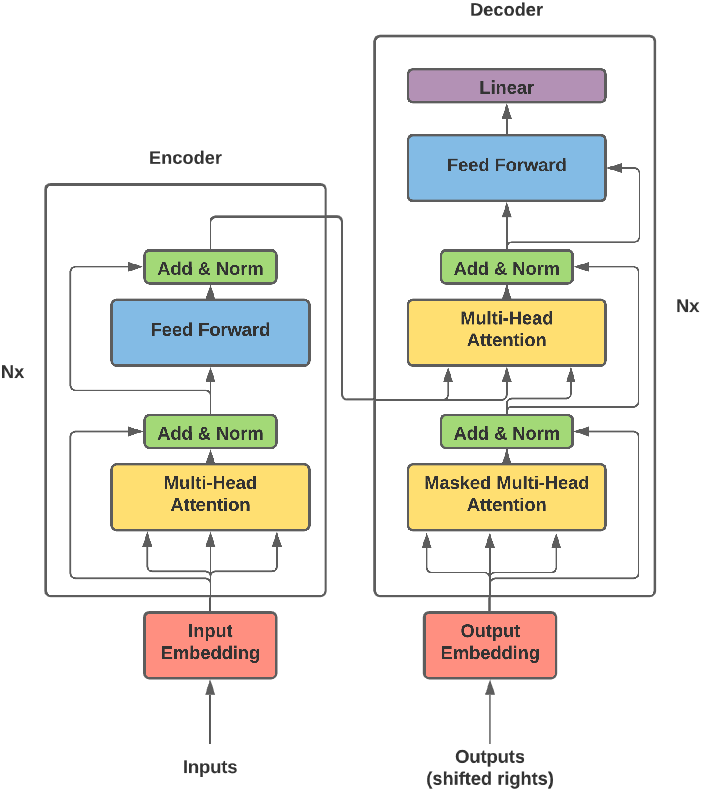
The MOT Model Architecture and Components.

## Results

### Evaluation of models performance

To assess the performance of the models, we used the traditional classification metrics: the accuracy 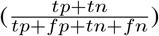, the Recall 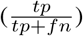, the Precision 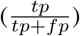 and the F1 score 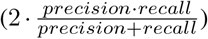. Since the dataset is imbalanced (see figure 3 in supplementary data), the F1 score is the metric used to asses the models performance. The MOT model is trained and evaluated on three partitions: a training set (70% of the dataset), a validation set (10% of the dataset) and a testing set (20% of the dataset). Table 1 provides a summary of the distribution of the examples in the dataset after the splitting before the data augmentation step. The MOT model metric scores are presented in the table 2 alongside with metric scores from OmiVAE (26), OmiEmbed (35), XOmiVAE (36) and Gene-Transfomer (37). OmiEmbed is an extension of OmiVAE that integrated a multi-task aspect to the original model previously introduced. It targets simultaneously three tasks: the classification of the tumour types (which is the main focus of this work), the regression (the age prediction and other clinical features) and the survival prediction. XOmiVAE is another extension of OmiVAE. It is an activation level-based interpretable deep learning models explaining novel clusters generated by VAE. GeneTransformer model is a transformer-based model combining a One-dimensional Convolutional-Neural Network (1D-CNN) and a transformer encoder block to extract features from 1D vectorized gene expression levels from TCGA samples. Thus it applies a DNN comprised of FCC to achieve the multi-classification task. Although the inputs of these models are not the same as the MOT model, since they all share the same prediction task i.e. the multiclassification of the 33 cancers of TCGA, we compare them. Indeed, OmiVAE, OmiEmbed and XOmiVAE used only 3 omics (miRNA, mRNA, and DNA methylation) without any missing omics views and GeneTransformer only one omic view (mRNA). Thus to make a fair comparison with MOT model, we evaluate the MOT model on 4 different tests set configuration: **(1)** on the samples with the 5 omics containing missing omics views, **(2)** only on the samples with the 3 omics (miRNA, mRNA, and DNA methylation) without missing omics views, **(3)** only the samples with only the mRNA omic and **(4)** on the samples with the 5 omics data without missing omics views. All results other than MOT are reported directly from their original article.

**Table 1.** Statistics distribution of the samples in the splitting of the dataset. The first part of the table give the statistic distribution of the missing omics views in the different part of the dataset. The second part shows the repartition of the different type of missing views.

|  | Train | Valid | Test |
| --- | --- | --- | --- |
|  | Train size: 8820 | Valid size: 981 | Test size: 2451 |
| Samples with at least ONE missing views | 4595(52.10%) | 472(48.11%) | 1260(51.41%) |
| Samples with ONE missing views | 2681(30.4%) | 278(28.34%) | 733(29.91%) |
| Samples with TWO missing views | 760(8.62%) | 75(7.65%) | 222(9.06%) |
| Samples with THREE missing views | 549(6.22%) | 52(5.30%) | 159(6.49%) |
| Samples with FOUR missing views | 605(6.86%) | 67(6.83%) | 146(5.96%) |
| Samples without missing views | 4225(47.90%) | 509(51.88%) | 1191(48.6%) |
| Samples with missing CpG sites | 1904 | 205 | 541 |
| Samples with missing miRNA | 1086 | 107 | 279 |
| Samples with missing RNA | 923 | 93 | 222 |
| Samples with missing CNV | 1021 | 100 | 294 |
| Samples with missing Protein | 3334 | 347 | 902 |

There are interesting observations to be drawn from the results presented at table 2. The comparison of the MOT model vs. the models OmiVAE, OmiEmbed and XOmiVAE shows that the MOT performs as well as those models and sometimes depending of the metrics even better. Indeed, MOT**(2)** achieves a F1 score of 97.33% which is slightly less than the OmiVAE (97.5%). But, MOT**(2)** (97.33%) performs better than OmiEmbed (96.83%) and outperformed XOmiVAE (90%). In the other comparison case between MOT and GeneTransformer, MOT achieved a better performance than GeneTransformer. MOT**(3)** has 96.54% of F1-score while GeneTransformer has 95.64%. We also evaluate the performance of the MOT model based on the availability of all the omics views in the samples. MOT**(4)** achieves a F1-score of 98.37% which is better than MOT**(1)** F1-score of 96.74%. This was the expected result, as most of the models tend to perform better when all the data are available. Table 5, in supplementary data, presents the classification report obtained with scikit-learn. Other than Rectum Adenocarcinoma (READ) cancer, MOT performs well on all remaining cancer. In table 7 in supplementary data, we also present the classification report for the experiment with all the views available for each sample.

**Table 2.**
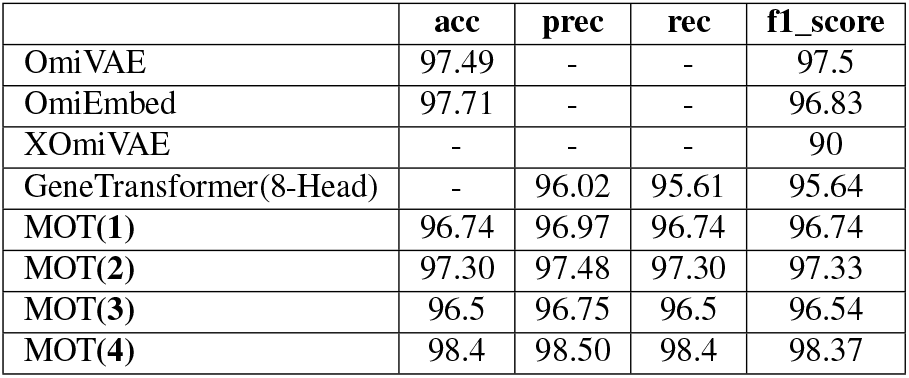
Performance metrics of the models. MOT is evaluated on the following settings: **(1)** on the samples with the 5 omics containing missing omics views, **(2)** only on the samples with the 3 omics (miRNA, mRNA, and DNA methylation) without missing omics views, **(3)** only the samples with the mRNA omic and **(4)** on the samples with the 5 omics data without missing omics views. The metrics performance results of OmiVAE, OmiEmbed, XOmiVAE and GeneTransformer are reported directly from their respective papers. ‘-’ means that metrics was not reported in their original papers.

**Table 3.** Omics views with the highest attention weights for each cancer.

| Cancers | CNVs | DNA methylation | miRNA | mRNA | protein |
| --- | --- | --- | --- | --- | --- |
| ACC |  | ✓ |  | ✓ |  |
| BLCA |  | ✓ |  | ✓ |  |
| BRCA |  |  | ✓ | ✓ |  |
| CESC |  | ✓ | ✓ | ✓ |  |
| CHOL |  | ✓ | ✓ | ✓ |  |
| COAD |  |  | ✓ | ✓ |  |
| DLBC |  | ✓ | ✓ | ✓ |  |
| ESCA |  | ✓ | ✓ | ✓ |  |
| GBM |  | ✓ | ✓ | ✓ | ✓ |
| HNSC |  | ✓ | ✓ | ✓ |  |
| KICH |  | ✓ | ✓ | ✓ |  |
| KIRC |  |  | ✓ | ✓ |  |
| KIRP |  | ✓ | ✓ | ✓ |  |
| LAML |  | ✓ | ✓ | ✓ |  |
| LGG |  | ✓ | ✓ | ✓ | ✓ |
| LIHC |  | ✓ | ✓ | ✓ |  |
| LUAD |  | ✓ | ✓ | ✓ |  |
| LUSC |  | ✓ | ✓ | ✓ |  |
| MESO |  | ✓ | ✓ | ✓ |  |
| OV | ✓ |  | ✓ | ✓ |  |
| PAAD |  | ✓ | ✓ | ✓ |  |
| PCPG |  | ✓ |  | ✓ |  |
| PRAD |  | ✓ |  | ✓ |  |
| READ |  |  | ✓ | ✓ |  |
| SARC |  | ✓ | ✓ | ✓ |  |
| SKCM |  | ✓ |  | ✓ |  |
| STAD |  | ✓ | ✓ | ✓ |  |
| TGCT |  | ✓ | ✓ | ✓ |  |
| THCA |  | ✓ | ✓ | ✓ |  |
| THYM |  | ✓ | ✓ | ✓ |  |
| UCEC |  | ✓ | ✓ | ✓ |  |
| UCS |  | ✓ | ✓ | ✓ |  |
| UVM |  | ✓ | ✓ | ✓ |  |

**Table 4.** Study Abbreviations

| Study Abbreviation | Study Name |
| --- | --- |
| ACC | Adrenocortical carcinoma |
| BLCA | Bladder Urothelial Carcinoma |
| BRCA | Breast invasive carcinoma |
| CESC | Cervical squamous cell carcinoma and endocervical adenocarcinoma |
| CHOL | Cholangiocarcinoma |
| COAD | Colon adenocarcinoma |
| DLBC | Lymphoid Neoplasm Diffuse Large B-cell Lymphoma |
| ESCA | Esophageal carcinoma |
| GBM | Glioblastoma multiforme |
| HNSC | Head and Neck squamous cell carcinoma |
| KICH | Kidney Chromophobe |
| KIRC | Kidney renal clear cell carcinoma |
| KIRP | Kidney renal papillary cell carcinoma |
| LAML | Acute Myeloid Leukemia |
| LGG | Brain Lower Grade Glioma |
| LIHC | Liver hepatocellular carcinoma |
| LUAD | Lung adenocarcinoma |
| LUSC | Lung squamous cell carcinoma |
| MESO | Mesothelioma |
| OV | Ovarian serous cystadenocarcinoma |
| PAAD | Pancreatic adenocarcinoma |
| PCPG | Pheochromocytoma and Paraganglioma |
| PRAD | Prostate adenocarcinoma |
| READ | Rectum adenocarcinoma |
| SARC | Sarcoma |
| SKCM | Skin Cutaneous Melanoma |
| STAD | Stomach adenocarcinoma |
| TGCT | Testicular Germ Cell Tumors |
| THYM | Thymoma |
| THCA | Thyroid carcinoma |
| UCEC | Uterine Corpus Endometrial Carcinoma |
| UCS | Uterine Carcinosarcoma |
| UVM | Uveal Melanoma |

**Table 5.** Classification performance of the MOT on each cancer label

|  | Cancers | precision | recall | f1-score | support |
| --- | --- | --- | --- | --- | --- |
|  | ACC | 0.96 | 0.92 | 0.94 | 24 |
|  | BLCA | 0.96 | 0.99 | 0.97 | 89 |
|  | BRCA | 1.00 | 1.00 | 1.00 | 255 |
|  | CESC | 0.96 | 0.91 | 0.93 | 55 |
|  | CHOL | 1.00 | 0.86 | 0.92 | 7 |
|  | COAD | 0.94 | 0.85 | 0.89 | 118 |
|  | DLBC | 1.00 | 1.00 | 1.00 | 7 |
|  | ESCA | 0.94 | 1.00 | 0.97 | 34 |
|  | GBM | 0.98 | 0.92 | 0.95 | 132 |
|  | HNSC | 1.00 | 0.98 | 0.99 | 111 |
|  | KICH | 0.96 | 1.00 | 0.98 | 22 |
|  | KIRC | 0.99 | 0.97 | 0.98 | 145 |
|  | KIRP | 0.93 | 0.97 | 0.95 | 71 |
|  | LAML | 0.89 | 0.97 | 0.93 | 40 |
|  | LGG | 0.98 | 1.00 | 0.99 | 105 |
|  | LIHC | 0.99 | 1.00 | 0.99 | 86 |
|  | LUAD | 0.95 | 0.95 | 0.95 | 128 |
|  | LUSC | 0.95 | 0.95 | 0.95 | 115 |
|  | MESO | 0.93 | 1.00 | 0.96 | 13 |
|  | OV | 0.95 | 0.96 | 0.96 | 122 |
|  | PAAD | 0.98 | 1.00 | 0.99 | 50 |
|  | PCPG | 1.00 | 0.97 | 0.99 | 37 |
|  | PRAD | 1.00 | 1.00 | 1.00 | 106 |
|  | READ | 0.51 | 0.75 | 0.61 | 24 |
|  | SARC | 0.97 | 0.98 | 0.97 | 58 |
|  | SKCM | 1.00 | 0.99 | 0.99 | 87 |
|  | STAD | 1.00 | 0.98 | 0.99 | 96 |
|  | TGCT | 1.00 | 1.00 | 1.00 | 24 |
|  | THCA | 0.99 | 1.00 | 1.00 | 113 |
|  | THYM | 1.00 | 1.00 | 1.00 | 22 |
|  | UCEC | 0.95 | 0.98 | 0.97 | 129 |
|  | UCS | 0.88 | 0.88 | 0.88 | 8 |
|  | UVM | 1.00 | 1.00 | 1.00 | 18 |
| **accuracy** |  |  |  | 0.97 | 2451 |
| **macro avg** |  | 0.96 | 0.96 | 0.96 | 2451 |
| **weighted avg** |  | 0.97 | 0.97 | 0.97 | 2451 |

**Table 6.** Classification performance of the MOT on each cancer label with all the 5 omics views available

|  | Cancers | precision | recall | f1-score | support |
| --- | --- | --- | --- | --- | --- |
|  | ACC | 1.00 | 1.00 | 1.00 | 9 |
|  | BLCA | 0.97 | 0.99 | 0.98 | 72 |
|  | BRCA | 1.00 | 1.00 | 1.00 | 125 |
|  | CESC | 1.00 | 0.89 | 0.94 | 27 |
|  | CHOL | 1.00 | 0.67 | 0.80 | 3 |
|  | COAD | 0.96 | 0.88 | 0.92 | 51 |
|  | DLBC | 1.00 | 1.00 | 1.00 | 6 |
|  | ESCA | 1.00 | 1.00 | 1.00 | 17 |
|  | GBM |  |  |  |  |
|  | HNSC | 1.00 | 1.00 | 1.00 | 60 |
|  | KICH | 1.00 | 1.00 | 1.00 | 15 |
|  | KIRC | 1.00 | 1.00 | 1.00 | 44 |
|  | KIRP | 1.00 | 1.00 | 1.00 | 44 |
|  | LAML |  |  |  |  |
|  | LGG | 1.00 | 1.00 | 1.00 | 82 |
|  | LIHC | 0.97 | 1.00 | 0.99 | 38 |
|  | LUAD | 0.97 | 0.98 | 0.98 | 64 |
|  | LUSC | 0.95 | 0.95 | 0.95 | 40 |
|  | MESO | 1.00 | 1.00 | 1.00 | 9 |
|  | OV |  |  |  |  |
|  | PAAD | 0.96 | 1.00 | 0.98 | 26 |
|  | PCPG | 1.00 | 0.95 | 0.97 | 19 |
|  | PRAD | 1.00 | 1.00 | 1.00 | 57 |
|  | READ | 0.54 | 0.78 | 0.64 | 9 |
|  | SARC | 1.00 | 1.00 | 1.00 | 46 |
|  | SKCM | 1.00 | 1.00 | 1.00 | 59 |
|  | STAD | 1.00 | 1.00 | 1.00 | 48 |
|  | TGCT | 1.00 | 1.00 | 1.00 | 15 |
|  | THCA | 1.00 | 1.00 | 1.00 | 68 |
|  | THYM | 1.00 | 1.00 | 1.00 | 18 |
|  | UCEC | 0.95 | 0.98 | 0.97 | 64 |
|  | UCS | 1.00 | 0.86 | 0.92 | 7 |
|  | UVM | 1.00 | 1.00 | 1.00 | 2 |
| accuracy |  |  |  | 0.98 | 1144 |
| macro avg |  | 0.98 | 0.96 | 0.97 | 1144 |
| weighted avg |  | 0.99 | 0.98 | 0.98 | 1144 |

**Table 7.** Best MOT model parameters

|  |  |
| --- | --- |
| data_size | 2000 |
| dataset_views_to_consider | all |
| exp_type | data_aug |
| activation | relu |
| batch_size | 256 |
| d_ff_enc_dec | 2048 |
| d_input_enc | 2000 |
| d_model_enc_dec | 512 |
| dropout | 0.44374742780410337 |
| early_stopping | True |
| loss | ce |
| lr | 0.00039893650505836597 |
| lr_scheduler | cosine_with_restarts |
| n_epochs | 500 |
| n_heads_enc_dec | 8 |
| n_layers_dec | 1 |
| n_layers_enc | 6 |
| nb_classes_dec | 33 |
| optimizer | Adam |
| weight_decay | 0.005744062413504335 |
| seed | 42 |
| class_weights | [4.03557312, 0.85154295, 0.30184775, 1.18997669, 8.25050505, 0.72372851, 7.73484848, 1.81996435, 0.62294082, 0.61468995, 4.07992008, 0.49969411, 1.07615283, 1.85636364, 0.7018388, 0.84765463, 0.60271547, 0.62398778, 4.26750261, 0.61878788, 1.89424861, 1.98541565, 0.65595888, 2.05123054, 1.37001006, 0.77509964, 0.76393565, 2.67102681, 0.64012539, 2.94660895, 0.64012539, 6.51355662, 4.64090909] |

### Macro interpretability

In the previous section, we demonstrated the model’s ability to predict accurately the various cancer types. Here, we further investigate the model ability to provide a level of interpretability. The aim of this analysis is to find which are the most important omics views and their individual impact on the model decision. In order to do this, an analysis of the multi-heads attention layers of the transformer model was performed. The goal is to investigate for each tumour the most impacting omics views on the decision output of the MOT model for this particular tumour. To do so, all the weights of all the layers are combined from each attention head. The weights are summed, the average is calculated, and a reduction is performed to obtain 5*5 arrays for each cancer sample. Then, these arrays are used to obtain heat maps of the interactions between all the omics views. We extract the omics views from those heat maps with the highest attention weights implying the most impact for each cancer. Table 3 presents the finding. Most of the attention weights are on the combination of the mRNA, the miRNA and the DNA methylation omics views. This is observed in 21 cancer cases. The second most observation is the focus of the attention weights on the combination of mRNA and DNA Methylation which occurs 4 times. In only two cases, we have an attention focus on 4 views: the Glioblastoma multiform (GBM) and Brain Lower Grade Glioma (LGG) cancers for which the model focus on the combination of mRNA, miRNA, DNA methylation and protein. The important information from this analysis is that the MOT model uses information from multiple omics views (mostly 3) instead of just focusing on a single one. Moreover, to analyze the impact of the omics views with the highest attention scores, for each cancer, the views identified in the table 3 are removed from the test set for each cancer, and MOT is re-evaluated. In figure 2 we illustrate the variation of the f1 scores. There is a degradation for all of the tumours when these omics are turned off. This observation supports the importance of these particular omics for the tumours.

**Fig. 2.**
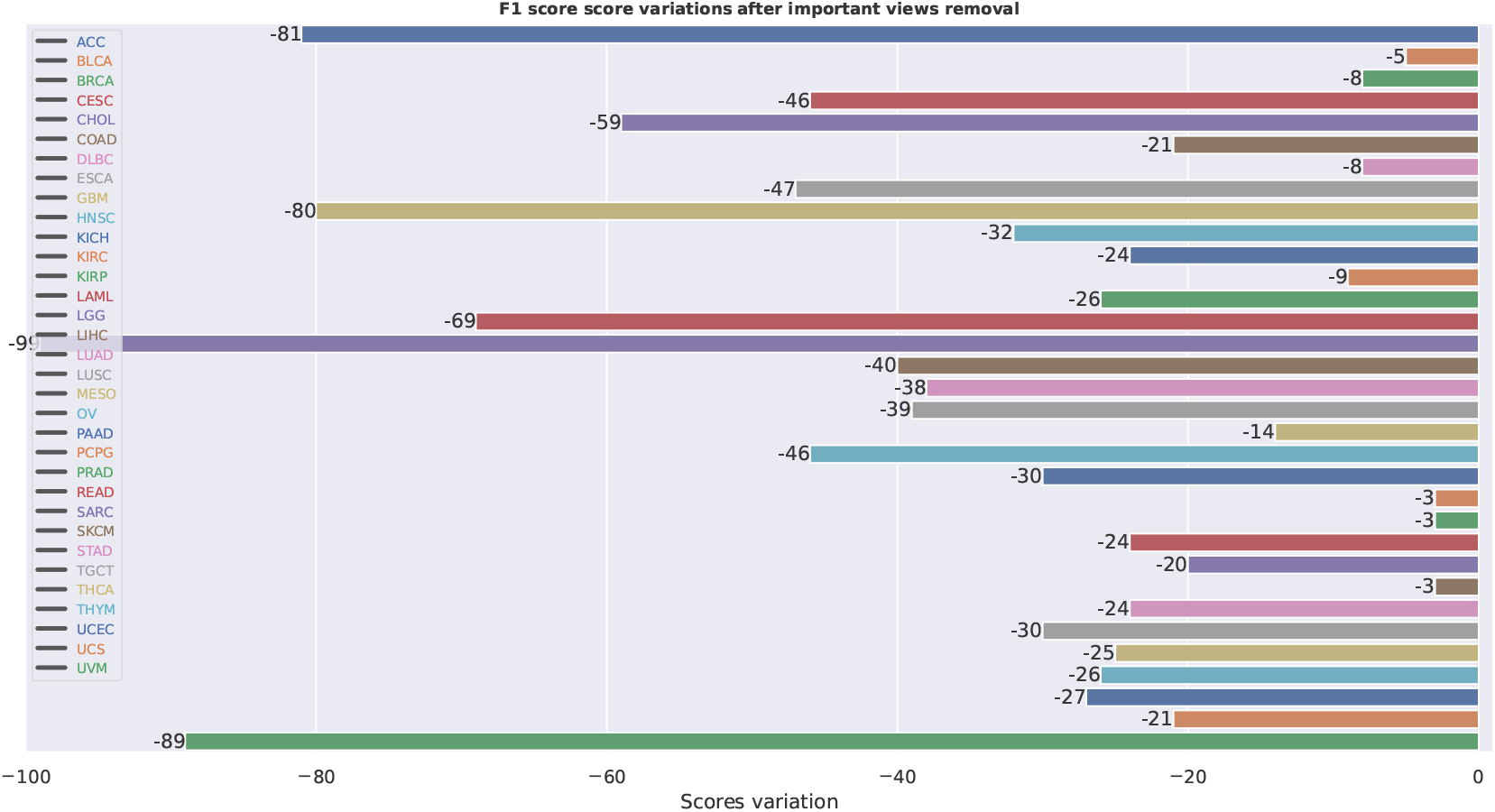
Metric evaluation of the MOT model for each cancer with each of views with the highest attention removed from the test set.

**Fig. 3.**
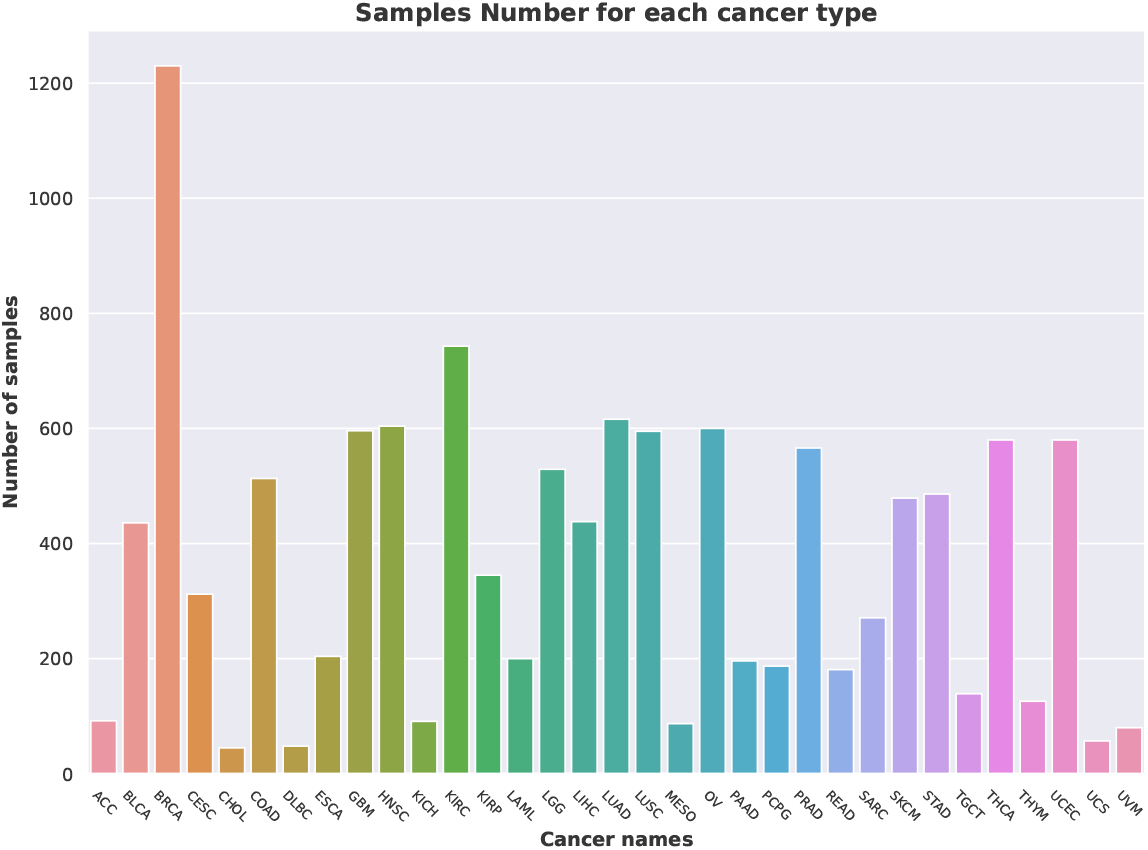
Distribution of the cancer in the dataset.

## Discussion and Conclusions

This paper introduces MOT: a multi-omics transformer for multiclass classification tumour types predictions. The model is based on a deep learning architecture, the transformer architecture with attention heads mechanisms (29). The scarcity of certain omics data makes multi-omic studies difficult and prevents using the full range of omics. Nevertheless, from the UCSC Xena data portal, five omics data type (CNVs, DNA-methylation, miRNA, mRNA and proteins) were extracted to build a multi-omics dataset. These omics data each have various feature space sizes ranging from a vast feature space (396066 original features for DNA methylation) to a relatively small feature space (259 original features for protein). This variation requires a quasi-mandatory preprocessing step to integrate the data correctly. These steps consist of a dataset dimension reduction via a feature selection and padding the missing views. The padding was done by replacing the values per 0, a bit drastic but our initial choice. After the preprocessing steps, the MOT model was trained and evaluated on the multi-omics dataset. The hyper-parameter optimization, a crucial step in machine learning problems, was done with Optuna (38), an open-source hyper-parameter optimization framework to automate hyper-parameter search. Through the training phase, a data augmentation step was performed. This step allows to diversify the type and the number of examples seen during the training phase with the primary purpose of increasing the model robustness. From the basic experiment scheme (i.e. train-test-validation scheme) the MOT model obtains a F1-score of 96.74% (see MOT**(1)** in table 2). Compared to other models presented in the table 2, the MOT model is not technically the best model. However, it does not use the same input data although they all have the same prediction task. In order to have a fair comparison of the MOT model, multiple evaluations were performed. We assessed the MOT performance on different test set: **(2)** only on the samples with the 3 omics (miRNA, mRNA, and DNA methylation) without missing omics views, **(3)** only the samples with the mRNA omic and **(4)** on the samples with the 5 omics data without missing omics views. The first evaluation on the samples with only 3 omics is to compare the model to the OmiVAE, OmiEmbed and XOmiVAE models. The performance reported in the table 2 demonstrate that MOT(**(2)**) are about the same or even better depending on the metrics. The second evaluation on the samples with the mRNA omic is to compare MOT to the GeneTransformer model. In this case, we can observe that the MOT performs better than the GeneTransformer. In this case, our model benefits from the contribution of the different omics views during the training phase. The last experiment was to show the performance of the model in the best-case scenario i.e. on the samples with the 5 omics data without missing omics views. In this case, the MOT model outperformed all the other experiments cases and other models with an F1-score of 98.37%. This demonstrates the excellent prediction capability of the MOT model under ideal conditions. It also emphasises the importance of using multi-omics data. To our knowledge, this is the first model able to integrate up to five omics views and be as efficient on the multiclass classification prediction task. The parameters of the best model obtained are presented in the table 7 in supplementary data.

The internal structure of the model, i.e. the attention mechanism heads, gives the MOT model a distinctive edge worth exploiting. The attention weights can help discover the most impactful views in the model decision process in general and for each cancer types. This identification will help the clinical decision-making process to better allocate resources to acquire certain specific omics views for certain tumour types. Table 3 shows the results of the analysis of the heatmaps of the attention weights. From this table we can draw the conclusion that the mRNA omic view is important for the prediction task no matter the tumour types. This omic view is followed by the DNA methylation which is the second omic view most weighted by the model and generally in combination with the mRNA omic view. This is followed by the miRNA omics views which is the 3rd most activated omic view. Another important observation from the table 3 is that at least 2 omics views are necessary for the prediction task and most of the time all the 3 principal omics (mRNA, DNA methylation and miRNA) are used. For only two tumours, GBM and LGG, the MOT model uses the protein omic view. This can be explained by the fact that this is the less developed omic view since not enough features are available and produced for this omic view. The lack of representation and probably the misrepresentation could lead the proteomic view to be less important in the decision-making process. The only case where the MOT model uses the CNVs omic view is for the Ovarian serous cystadenocarcinoma (OV) cancer. To corroborate these findings, we elected to test the MOT model on a subset: the same test set at least samples wise but without the most impactful views determined by the model for each cancer and presented in the table 3. The goal is to demonstrate the impact of those views on performance degradation. Mixed results are obtained (see figure 2). As expected, all the performance decreases when the most impactful omics views per cancer are removed from the test set. The multi-omic transformer model introduced here covers many important areas of multi-omics studies. Although cancer has historically been viewed as a disorder of proliferation, recent evidence has suggested that it should also be considered, in part, a metabolic disease (39–42). Thus, we wonder if the mRNA importance observed here is not due to an over-representation. To ensure a better understanding of the complex phenomena which is cancer, the possible next steps of this model is to integrate the metabolomic view into the fold. This would imply a different integration process and a more comprehensive picture of the cancer disease.

## Ethics approval and consent to participate

Not applicable

## Consent for publication

Not applicable

## Availability of data and materials

The datasets generated and/or analysed during the current study are available in the Xena Data portal repository, cohort: TCGA Pan-Cancer (PANCAN): https://pancanatlas.xenahubs.net. The CNVs data retrieved are available in the Synapse database, accession number: syn5011220.1. The DNA methylation data retrieved are available in the Synapse database, accession number: syn4557906.9. The mRNA data retrieved are available in the Synapse database, accession number: syn4976369.3. The miRNA data retrieved are available in the Synapse database, accession numbers: syn6171109 and syn7201053. The proteins data retrieved are available in the Synapse database, accession number: syn4216793.3. The code is available at: https://github.com/dizam92/multiomic_predictions.

## Authors’ contributions

MAO and PT conceived the experiment(s). MAO conducted the experiment(s). MAO, PT, JC and FL analyzed the results, and wrote the manuscript. All authors reviewed the manuscript. All authors read and approved the final manuscript.

## Acknowledgements

A special thanks to Rogia Kpanou for her inputs in this work. We also acknowledge the support of Compute Canada for providing additional computational support and also Dr Jacques Corbeil’s Canada Research Chair in Medical Genomics.

## Supplementary Note 1: Supplementary Tables and Figures

